# TMS bursts can modulate local and networks oscillations during lower-limb movement

**DOI:** 10.1101/2020.08.19.257980

**Authors:** Arturo I. Espinoza, Jamie L. Scholl, Arun Singh

**Author notes:** These authors contributed equally. **Correspondence to:** Dr. Arun Singh, Division of Basic Biological Sciences, Sanford School of Medicine, University of South Dakota, 414 E. Clark St. Vermillion, SD, 57069, USA.

## Abstract

**Introduction:** Lower-limb motor functions involve processing information via both motor and cognitive control networks. Measuring oscillations is a key element in communication within and between cortical networks during high order motor functions. Increased midfrontal theta oscillations are related to improved lower-limb motor performances in patients with movement disorders. Non-invasive neuromodulation approaches have not been explored extensively to understand the oscillatory mechanism of lower-limb motor functions. This study aims to examine the effects of repetitive transcranial magnetic stimulation (rTMS) on local and network EEG oscillations in healthy elderly subjects.

**Methods:** Eleven healthy elder subjects (67-73 years) were recruited via advertisements, and underwent both active and sham stimulation procedures in a random, counterbalanced design. TMS bursts (θ-TMS; 4 pulses/sec) were applied over the midfrontal lead (vertex) before a GO-Cue pedaling task, and signals were analyzed using time-frequency methods.

**Results:** TMS bursts increase the theta activity in the local (p=0.02), as well as the associated network during the lower-limb pedaling task (p = 0.02). Furthermore, after task-related TMS burst sessions, increased resting-state alpha activity was observed in the midfrontal region (p= 0.01).

**Conclusion:** Our study suggests the ability of midfrontal TMS bursts to directly modulate local and network oscillations in a frequency manner during lower-limb motor task. TMS burst-induced modulation may provide insights into the functional roles of oscillatory activity during lower-limb movement in normal and disease conditions.

## INTRODUCTION

Increasing evidence has shown the role of cortical oscillations in conveying information within motor and cognitive control networks.^1–3^ Within the motor and cognitive research field, midfrontal theta, beta, and gamma oscillations have been associated with motor tasks with cognitive recruitment such as gait or lower-limb movements.^4–6^ Cognitive contributions to lower-limb movement in elderly people (>50 years old) and patients with movement disorders are well-acknowledged in movement sciences.^7,8^ Additionally, the role of midfrontal theta oscillations in cognitive control has been studied extensively in neuropsychiatric and movement disorders patients.^3,9^ Animal studies have shown increased theta oscillations around the anterior cingulate cortex and frontal cortex, similar to frontal midline theta oscillations in humans, during attentional motor tasks.^10,11^ These studies have spurred a growing interest in the manipulation of midfrontal theta oscillations to improve attentional motor tasks. A better understanding of their causal physiological role can be achieved in the coding of movements involving attentional processes. Indeed, this endeavor has the potential to alleviate lower-limb motor problems in patients with movement disorders.

Non-invasive cortical stimulation techniques such as transcranial magnetic and electrical stimulation (TMS and tES) have emerged as promising neuromodulation methods, enabling direct modulation of oscillations in the local and related networks.^12,13^ TMS, specifically repetitive TMS (rTMS), appears to be a particularly effective neuromodulation tool to target spatially confined cortical regions, and in modulating oscillations in the local region and associated neural networks.^14,15^ In comparison to a single pulse, TMS bursts can induce long-term plastic changes in the motor and cognitive networks and modulate neuronal oscillations effectively, leading to an increased interest in therapeutic possibilities.^16,17^ Recent findings have shown that TMS bursts in the motor cortical region can improve gait and balance problems in Parkinson’s Disease (PD) patients^18,19^, however, the mechanisms by which TMS bursts exert neuromodulatory effects on lower-limb motor performance remain unknown.

TMS’s ability to modulate motor networks in combination with EEG’s ability to record changes in oscillatory network activity with high temporal resolution makes it an ideal method to understand underlying motor performance.^20,21^ Nevertheless, it remains unclear whether TMS bursts (θ-TMS; 4 arrhythmic pulses in a second) result in entrainment for theta frequency bands in the local stimulated brain region. Furthermore, it is not known how locally modulated oscillations impact resultant activity in the associated cortical regions during lower-limb movement in an elderly population. To address these questions, we analyzed EEG recordings in healthy elderly subjects, while applying TMS bursts to the midfrontal region (at vertex). The motor and cognitive deficits thought to be related to theta oscillations are typically found in a patient population such as PD.^3,6^ As the average age of initial onset for PD is 60, baseline network activity was determined in a non-patient population within a similar age group, to better understand how these networks are affected in a comparable control group, prior to examining them in a patient population with network dysfunction.

## METHODS

### PARTICIPANTS

Eleven healthy elderly subjects (mean ± sem; age 70 ± 3 years; 6F/5M) were recruited in Iowa via advertisements to participate in the study. All protocols were approved by the University of South Dakota Institutional Review Board and the University of Iowa Institutional Review Board in accordance with the Declaration of Helsinki. Recruitment was limited to older adults, as a healthy, age-appropriate control group for future PD patient population studies. All subjects were free from orthopedic conditions, and were screened for exclusionary criteria, using our laboratory’s standard questionnaire, which consists of neurological and mental health history, psychoactive medication use, presence of a cardiac pacemaker or any other metal implant. All participants provided written informed consent to participate in the study.

### LOWER-LIMB PEDALING MOTOR TASK AND ANALYSIS

Similar to our previous study,^6^ a pedaling motor task was used to study lower-limb motor control. Participants were instructed to start pedaling upon seeing the GO-cue and to stop the pedals in the starting position after one rotation until the next GO-cue. This type of motor task induces minimal movement-related artifact and requires bilateral lower-limb coordination similar to gait. Furthermore, this paradigm can be used to investigate feedback-based neuromodulation. The pedaling task involves 2 blocks, each consisting of 25 trials during which the subject remains seated. Fifty total trials provide adequate signal-to-noise ratios for subsequent EEG analysis, and also prevents fatigue in elderly participants as demonstrated in our EEG work.^6,22^ For each trial, a warning cue briefly appeared on a stimulus presentation computer screen. Afterwards, a TMS burst was delivered to the midfrontal region (Cz) before the GO-Cue, triggered by TTL input. The 2 s GO-Cue appeared 3 ± 0.6 s after the 4^th^ TMS pulse discharge (Fig. 1A), with an inter-trial interval of 3 ± 0.1 s.

**FIG. 1.**
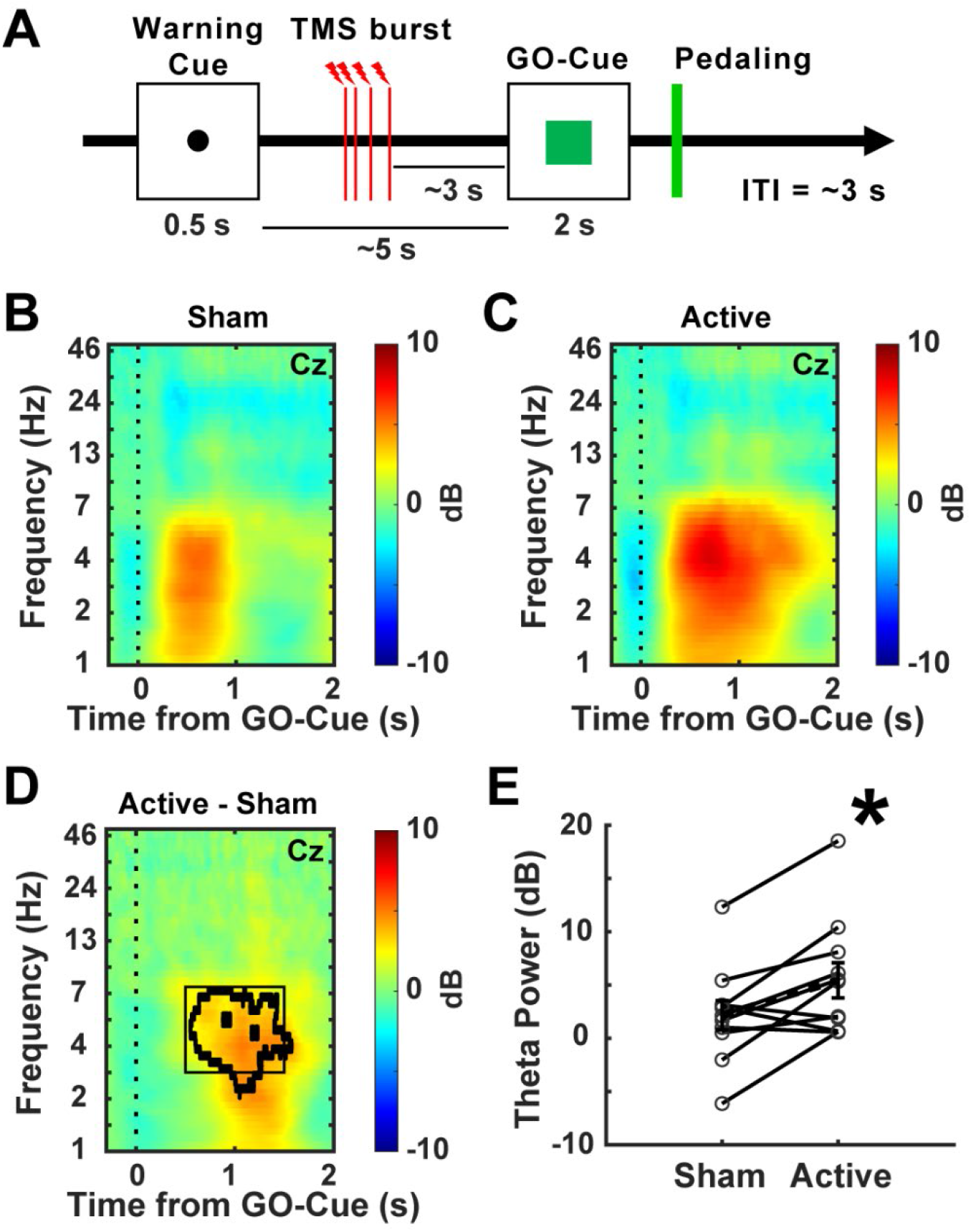
θ-TMS bursts increased pedaling task-related theta-band activity in the midfrontal region. A) Schematic diagram of the trial of EEG-TMS pedaling task. B-D) Average time-frequency analysis across subjects showed increased theta-band (3-7 Hz) power at midfrontal scalp electrode (vertex or “Cz”) during pedaling task after active TMS bursts as compared to sham in healthy elderly subjects. E) Plot displayed the mean power values in tf-ROIs (theta-band power value under black box) during pedaling task. D: p<0.05 outlined in bold lines. E: *p<0.05 vs sham stimulation; dashed line with error bar represents the mean and standard error of mean.

The total duration of each pedaling trial averaged 10.4 ± 0.7 s across subjects. A 3-axis accelerometer was attached to the subject’s left ankle. Afterwards, all three acceleration signals were averaged to generate a single signal. Linear speed for each trial was analyzed in single signal during the 0-2 s time window following the GO-Cue signal, as participants took approximately 2 s to complete one pedal rotation. Signals were detrended, and a 5 Hz low-pass filter was applied prior to computing the linear speed. Mean linear speed for each subject (m/s) was calculated.

### ACTIVE AND SHAM TMS TO THE MIDFRONTAL REGION

We used a figure-of-eight TMS coil (MC-B70) connected to a MagVenture (MagVenture, Inc., Alpharetta GA) Stimulator (MagPro X100) in both active and sham stimulation conditions over the midfrontal (“Cz” vertex) region according to the EEG 10/20 electrode position. We selected the midfrontal location for the TMS to modulate theta oscillations in the target and connected regions.^3,6^ Previous studies have demonstrated that stimulation over the vertex can increase leg motor cortex excitability bilaterally, and induce the most stable responses in the leg motor area.^23,24^ The current study followed the suggested TMS methodological checklist to improve the quality of data collection and reporting results.^25^ For sham stimulation, the figure-of-eight coil was inverted relative to the target location and off-targeted, which reduced the magnetic field significantly to below the threshold necessary to modulate oscillations.^26,27^ Stimulation intensity was set at 90% of the individuals resting motor threshold (RMT). RMT was determined on each subject, and is the minimum intensity that elicited at least five finger twitches in response to ten consecutive single-pulses applied to the motor hot spot.

Participants were blinded to stimulation, and both active and sham stimulation conditions were counterbalanced and randomized, with a minimum of 60 min between sessions; which has been shown to exceed the response time for the modulatory effects of theta burst TMS on cognitive tasks.^28^ A single TMS burst (θ-TMS) was applied, which consisted of 4 consecutive arrhythmic biphasic pulses within 1 s, and onset of TMS pulses differed for each trial and participant.

### EEG-TMS RECORDING AND PREPROCESSING

Scalp EEG-TMS signals were collected during the lower-limb pedaling task using a 64-channel EEG actiCAP (Brain Products GmbH, Gilching, Germany) and pycorder software with high-pass anti-aliasing FIR filter of 0.1 Hz with order 1 and 10kHz sampling frequency. EEG signals were also collected for 2-3 min during the eyes-open resting-state condition after completing pedaling task.

Reference and grounded channels were electrodes Pz and Fpz, respectively. Channels that were considered unreliable due to muscle artifacts or white noise (Fp1, Fp2, FT10, TP9, and TP10) were removed. EEG signals were epoched around the GO-Cue (−0.6 s to 2.4 s) to exclude TMS-evoked artifacts occurring prior to GO-Cue. Epochs with excessively noisy EEG or muscle artifacts were excluded using a conjunction of the Fully Automated Statistical Thresholding for EEG artifact Rejection (FASTER) algorithm with default parameters^29^ and pop_rejchan from EEGLAB.^30^ Eye movement artifacts were removed using an automatic EEG artifact detection algorithm based on the joint use of spatial and temporal features (ADJUST) with default parameters.^31^ Subsequently, data was rereferenced to common average reference.

### EEG POSTPROCESSING ANALYSIS

Preprocessed (epoched) signals were used for time-frequency analysis by complex Morlet wavelets, as explained previously.^3,6,32^ Further, epochs were cut in –0.5 to +2.0 s windows, frequency bands between 1 and 50 Hz in logarithmically-spaced bins were chosen, and power was normalized by conversion to a decibel (dB) scale. Similar to prior studies, the baseline for each frequency consisted of the average power from –0.3 to –0.2 s prior to the onset of the GO-Cue.^6,32^ Time-frequency analyses was limited to electrode Cz, and a-priori time-frequency Regions of Interest (tf-ROI). tf-ROIs were preselected for the theta-band (3–7 Hz) and the time window of interest was +0.5 to +1.5 s following the GO-Cue. tf-ROIs mean power values were used for statistical comparisons, with the main analysis restricted on a-priori tf-ROI as it represents the specificity, and tf-ROIs topographic map was plotted and compared between active versus sham conditions.

Preprocessed signals were also used for time-frequency spectral coherence (magnitude-squared coherence) analysis. A –0.3 to –0.2 s window was used as a baseline to compute change in coherence, then spectral coherence algorithms were implemented as explained in detail by Cohen.^33^ Spectral coherence over trials between Fz and Cz (fronto-cortical network) and C3 and C4 (inter-hemispheric motor cortical network) was computed. The same priori tf-ROIs were used (as above) to export coherence values to compare active versus sham stimulation data.

EEG resting-state signals were preprocessed (similar to above) and Welch’s power spectral density estimate (pwelch) Matlab function was applied on epoched (3.5 s) signals to compute spectral properties. Relative power at delta and alpha frequency bands was quantified compared to the power over 0.1–50 Hz. The resting-state mean relative power, after the task-related active and sham TMS bursts, was compared. Resting-state analysis was focused on the Cz electrode.

Wilcoxon signed rank tests were used to compare behavior and electrophysiological data between active TMS and sham. Effect sizes were computed as Cohen’s d. Statistical analysis was focused on our a-priori hypothesis to determine the specificity and restricted to midfrontal (“Cz”) electrode and theta-band activity only. In addition to this, we employed threshold-based correction on the full time-frequency plots in order to reveal any other reliable differences between active and sham TMS, and these differences are shown as black contour in time-frequency plots. The alpha level was 0.05 and Bonferroni adjustments were applied with adjusted alpha level 0.026 where appropriate.

## RESULTS

Active TMS bursts (θ-TMS) significantly increased the theta-band power when compared to a within-subjects sham stimulation during pedaling task in healthy elderly subjects (z = −2.31, p = 0.02, Cohen’s d = −0.934, Fig. 1B–E, see Fig. S1 for each subject). Time-frequency plots show no changes in the other frequency bands power (Fig. 1B–E). Our results distinctly demonstrate that θ-TMS bursts (4 pulses/sec) can enhance the power in the burst frequency which is the theta range in our study.

Spectral coherence analysis between frontal leads (Fz-Cz) showed increased pedaling-task related coherence within the theta frequency range after active TMS (Fig. 2A–C, p<0.05, see the black contour in figure 2C). A significant increase in coherence in the tf-ROI theta-band was observed after active θ-TMS bursts as compared to sham (z = −2.2, p = 0.026, Cohen’s d = −0.70, Fig. 2D). Furthermore, we analyzed spectral coherence between left and right motor cortical leads (C3-C4) to observe coherent activity in the inter-hemispheric motor cortical network. Inter-hemispheric motor cortical network showed increased pedaling-task related coherence within the theta frequency range after active θ-TMS bursts (Fig. S2A–C; p<0.05, see the black contour). A significant increase in coherence in the theta-band was also seen in the inter-hemispheric motor cortical network after active θ-TMS bursts as compared to sham (z = −2.04, p = 0.04, Cohen’s d =-0.65, Fig. S2D). In line with previous reports, our results also suggest that TMS bursts may modulate the cortical network oscillations during lower-limb motor task.^15,34^

**FIG. 2.**
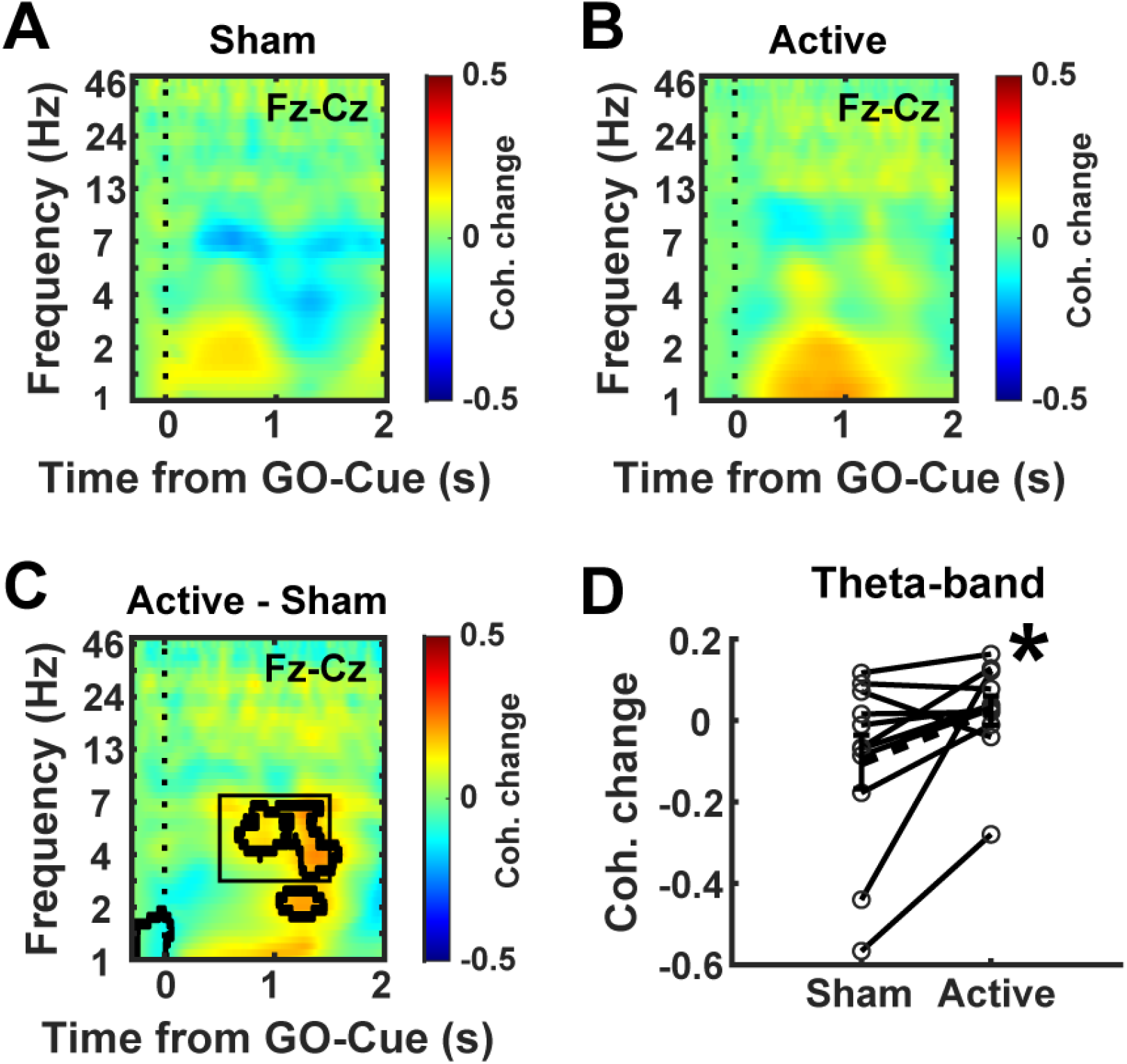
Spectral coherence in the fronto-cortical network was increased within theta-band. θ-TMS bursts can increase spectral coherence in the fronto-cortical network (Fz-Cz) in the theta frequency band during pedaling task in healthy elderly subjects. A) Sham and B) active TMS bursts induced changes in coherence from baseline during pedaling task. C) Difference in coherence between active and sham stimulation. D) Change in coherence from the baseline in the tf-ROIs: theta frequency band during active and sham TMS bursts. C: p<0.05 outlined in bold lines and black box represents the tf-ROIs in the theta frequency range. D: *p <0.05; dashed line with error bar represents the mean and standard error of mean.

Further, we analyzed poststimulation resting state signals to investigate the overall changes in the spectral properties after the active and sham θ-TMS burst sessions (Fig. 3A). No significant difference in relative power was found in the theta-band in the midfrontal lead after active TMS burst sessions compared to sham (z = 1.96, p = 0.05, Cohen’s d = 0.66, Fig. 3B). However, we found decrease in relative power in the alpha-band (z = 2.58, p = 0.01, Cohen’s d = 0.91, Fig. 3C) in the midfrontal lead. In line with current results, previous studies have also reported that following transcranial stimulation protocols, changes in resting-state alpha oscillations have been shown to modulate cognitive functions.^20,35,36^.

**FIG. 3.**
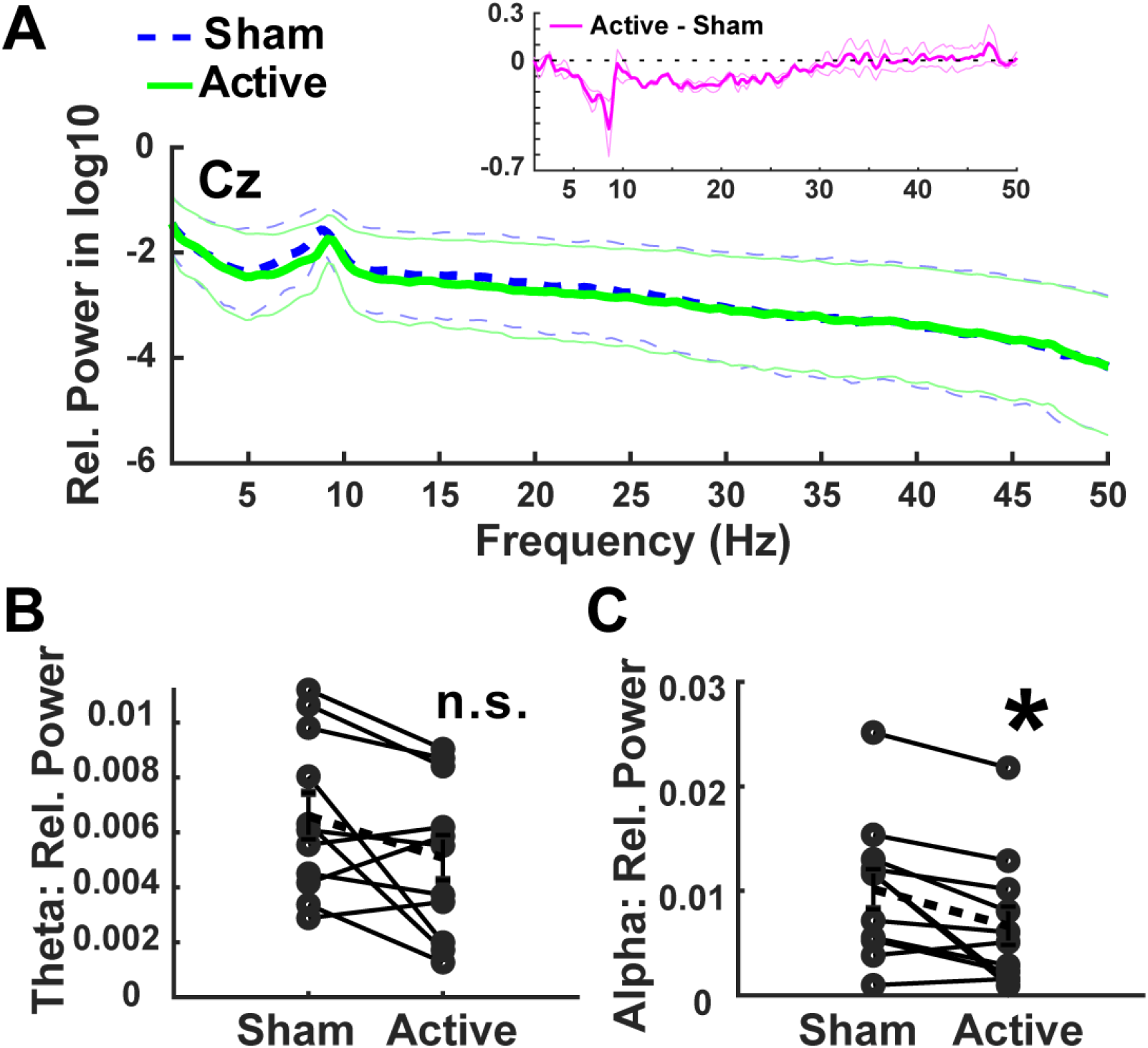
θ-TMS bursts sessions increased poststimulation resting state midfrontal alpha-band activity. A) Resting-state spectra after task-related active and sham θ-TMS bursts sessions. B-C) Graphs show no difference in theta activity but significant decreases in alpha activity after active TMS bursts sessions. a: thin dashed and solid lines represent the standard error of mean for sham and active stimulation, respectively. *p<0.05 vs. sham; dashed line with error bar represents the mean and standard error of mean; n.s. = non-significant.

Our data showed an increase in acceleration (Fig. 4B) and a time-dependent increase in pedaling speed following active θ-TMS bursts when compared to sham (Fig. 4C). However, we did not observe significant change in the pedaling speed in our subjects across trials (mean ± sem speed: sham = 0.2 ± 0.01 m/s; active = 0.3 ± 0.04 m/s; z = 0.8, p = 0.42, Cohen’s d = −0.23).

**FIG. 4.**
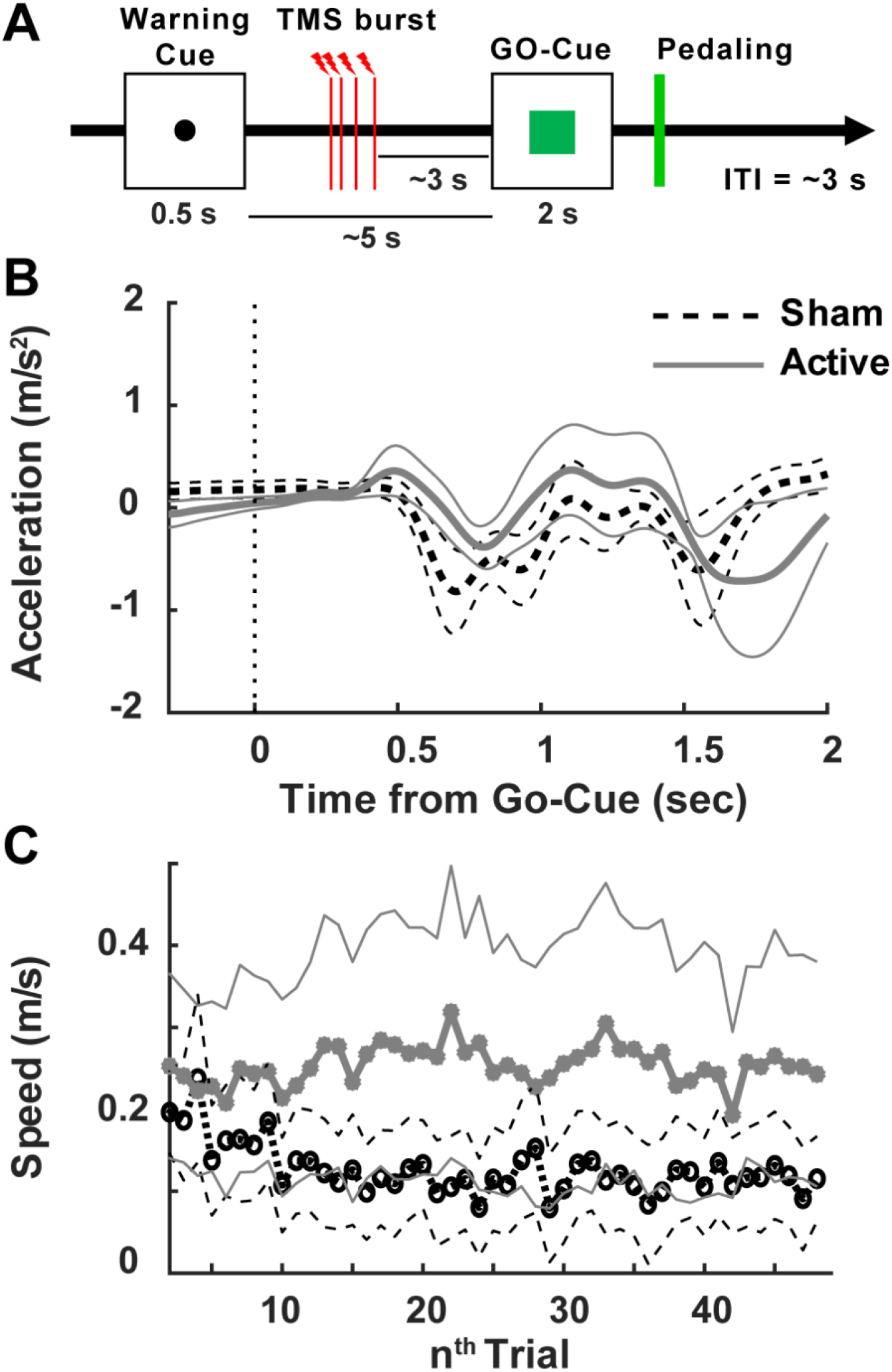
Effect of θ-TMS bursts on the lower-limb pedaling task. A) Mean acceleration signal after active and sham TMS bursts. B) Time-dependent changes in pedaling speed after active and sham TMS bursts. Thin dashed and solid lines represent the standard error of mean for sham and active stimulation, respectively.

## DISCUSSION

To investigate the immediate effects of TMS bursts on a lower-limb motor task, it is crucial to identify the actions of TMS on local and network oscillations. Our results suggest that TMS can modulate brain oscillations in direct interaction with the underlying generator, via synchronized midfrontal lower-limb task-related theta activity when targeting the theta (4-7 Hz) oscillations generator. We also demonstrate that θ-TMS bursts can enhance theta oscillations in the associated network. Results suggest that θ-TMS bursts may enhance the naturally occurring theta oscillations rather than impose artificial oscillations. The overall data support the hypothesis that TMS bursts may drive natural brain oscillations by entrainment, and this might be a possible mechanism via which TMS may act on the local and associated oscillatory activities to influence motor behavior.^20,34,37^

Movement disorder patients with lower-extremity dysfunctions have shown impaired cognitive control and decreased frontal theta oscillations, and thus show impaired kinematics of lower-limb movement ^6^. Attention is an important factor when performing lower-limb movements, especially when walking under dual-task conditions.^8,38^ Midfrontal theta oscillations may play the primary role in providing strong cognitive engagement during dual-tasks,^6,39^ contributing to the sensorimotor integration needed to perform lower-limb motor tasks. The current task-related EEG data reveal that θ-TMS bursts can upregulate targeted theta oscillations in higher-order frontal regions of the attention network and may enhance lower-limb movements.

Frontal beta oscillations also play a critical role in the top-down pathway conveying the information from preparatory and execution plans during lower-limb motor tasks, and these processes may malfunction in PD patients and other movement disorders.^2,40^ Increased beta-band power in the cortical and basal ganglia networks are associated with motor abnormalities in patients with PD.^2,41^ The dopamine precursor, levodopa, has shown the modulatory effects on the cortical and sub-cortical beta oscillations as well as the degree of motor impairment in patients with PD.^42,43^ The existence of two principal modes of synchronized oscillations within the human subthalamo-pallidal-thalamo-cortical circuit, at <30 Hz has been suggested in PD, and these oscillations are systematically modulated by motor task, thereby suggesting a functional role in movement.^44^ It has been shown that invasive deep brain stimulation (DBS) attenuates the beta-band activity and improves upper-limb movement in PD patients.^45,46^ However, fewer DBS studies have shown a therapeutic option for improving lower-limb movements such as gait in patients with PD.^47,48^ Similar to invasive studies, non-invasive TMS bursts approach may decrease cortical beta-band power and improve gait or lower-limb performances in PD.^19^ The overall data may support our previous hypothesis that increased frontal theta activity and reduced beta activity are associated with improvement in lower-limb movement in PD patients with gait abnormalities.^6^

High order lower-limb movements are a network phenomenon; and aberrations in the cortico-cortical and motor cortical networks have been observed in PD patients with gait abnormalities.^2,49^ Non-invasive TMS protocols can modulate oscillations in the target and associated networks during motor and cognitive tasks.^19,50^ In line with these studies, current results demonstrate the capacity of TMS bursts protocol to modulate theta oscillations in the cortico-cortical and motor cortical networks, which may be correlated with changes in the motor behavior even with the cognitive load. While published reports indicate that TMS bursts can improve motor performance in patients with movement disorders,^19,51^ we did not see a change in motor function after stimulation. Most likely, absent motor findings in our study reflect the low elderly subject numbers. However, uncoupling of EEG and motor effects of repetitive TMS is also a possibility.

Additionally, results show that θ-TMS bursts did not modulate resting-state theta oscillations, suggesting the change in theta activity is most-likely associated with GO-Cue-triggered motor performance.^3,6^ However, an increase in resting-state alpha-band activity following TMS bursts sessions demonstrates the overall poststimulation effect in the target region, which may also be related to improvement in motor and cognitive control systems.^20,36^ Notably, changes in alpha oscillations have been observed after transcranial neuromodulation in previous human studies,^35,52^ and these oscillations have been implicated in top-down cortical processing.^53^

The limitations of this study include: 1) Behavioral non-responders in the pedaling task, perhaps due to small participant numbers or the application of 90% RMT being insufficient to induce changes in pedaling speed, however, strong modulation in the theta-band activity was observed; 2) TMS was applied over the midfrontal region rather than specifically over the leg motor cortical area, however, current methods have been shown to induce leg motor cortex excitability in both hemispheres;^23,24^ In addition, TMS over midfrontal region can improve cognitive processing; 3) Use of a small, non-patient target population to explore the underlying mechanisms in the control of lower-limb motor networks prior to testing in a patient population rather than including the patient population in the experimental design. Nonetheless, data generated with this group will serve as a control for the age range in the patient population that will be examined in future studies.

## CONCLUSIONS

The current study may provide an efficient, non-invasive method to normalize brain oscillations that are often abnormal in conditions with motor disabilities. Specifically, by combining the TMS bursts protocol utilized with specific motor paradigms, this approach has the potential to investigate the causal links between midfrontal brain oscillations and lower-limb movement in humans.^6^ Further, this method may provide an understanding of the neurophysiological basis of modulations in the local and associated oscillatory networks, potentially contributing to novel rehabilitation regimens in patient populations such as PD.

## ACKNOWLEDGEMENTS

The authors thank all participants in this study. AIE is supported by the University of Iowa, IA. JLS and AS are supported by the Division of Basic Biomedical Sciences, University of South Dakota, Vermillion, SD, USA. We also thank Dr. Lee A. Baugh for his efforts towards improving our manuscript.

## SUPPLEMENTARY INFORMATION

### SUPPLEMENTARY FIGURES

**FIG. S1.**
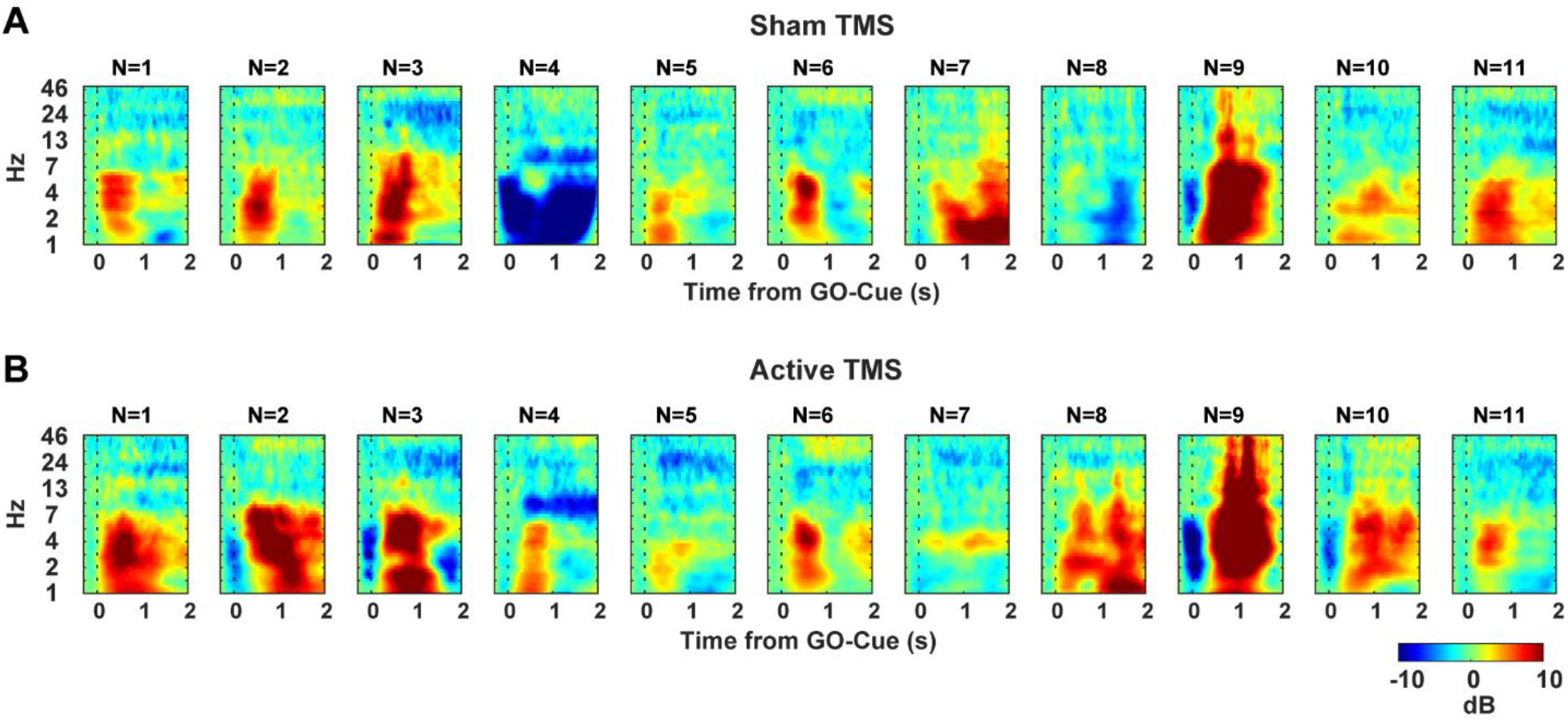
Active and sham θ-TMS bursts induced midfrontal activity in each subject. a-b) Pedaling-task related time-frequency plots in the midfrontal lead (Cz) for each healthy elderly subject after sham (A) and active (B) TMS bursts stimulations.

**FIG. S2.**
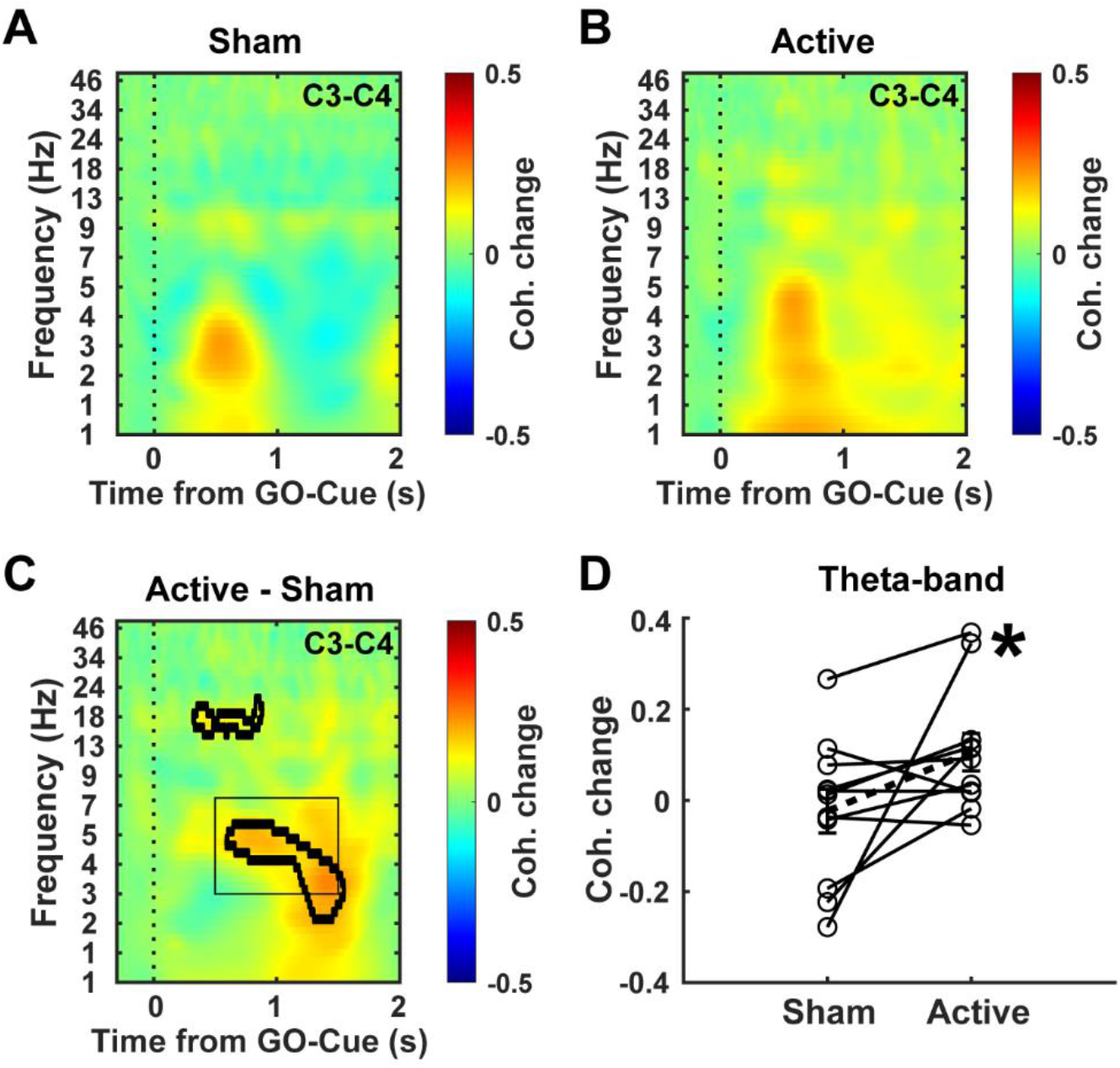
Spectral coherence in the inter-hemispheric motor cortical network was increased within theta-band. θ-TMS bursts can increase spectral coherence in the inter-hemispheric motor cortical network (C3-C4) within the theta frequency band during pedaling task in healthy elderly subjects. A) Sham and B) active TMS bursts induced changes in coherence from baseline during pedaling task. C) Difference in coherence between active and sham stimulation. D) Change in coherence from the baseline in the tf-ROIs: theta frequency band during active and sham TMS bursts. c: p<0.05 outlined in bold lines and black box represents the tf-ROIs in the theta frequency range. d: *p<0.05 vs. sham; dashed line with error bar represents the mean and standard error of mean.

